# Differential growth is a critical determinant of zebrafish pigment pattern formation

**DOI:** 10.1101/2021.06.11.448058

**Authors:** Jennifer P. Owen, Christian A. Yates, Robert N. Kelsh

## Abstract

The skin patterns of vertebrates are formed by complex interactions between pigment-producing cells during development. Adult zebrafish (*Danio rerio*), a model organism for investigating the underlying patterning processes, display alternating horizontal blue and golden stripes, generated by the self-organisation of three pigment cell-types. Mathematical studies in which these cells are modelled as individual agents communicating via short- and long-range interactions have produced breakthroughs in the understanding of pattern development. These models, incorporating all experimentally evidenced cell-cell interactions, replicate many aspects of wild-type and mutant zebrafish patterns. Although received wisdom suggested that initial iridophore distribution was pivotal in orienting patterning, here we show that growth can override its influence. Altered growth sequences can generate further pattern modulation, including vertical stripes and maze-like patterns. We demonstrate that ventrally-biased (asymmetric) growth of the skin field explains two key zebrafish pattern development features which are otherwise obscure (dorso-ventral pattern asymmetry, and predominant ventral-to-dorsal migration of melanophores) in wild-type and multiple zebrafish mutants, and in the related species *Danio nigrofasciatus*. By identifying biased growth as a novel patterning mechanism, our study will inform future investigations into the mechanisms and evolution of fish pigment patterning and vertebrate pigment pattern formation. Furthermore, our work has implications for the mechanistic basis of human pigmentation defects.

## Introduction

The diversity of skin pigment patterns in zebrafish and related *Danio* species, combined with a wealth of experimental observations, make the zebrafish a paradigmatic example of pigment pattern formation. Zebrafish pigment pattern formation involves diverse cell-cell interactions, which generate both WT and mutant pigment patterns^1-11^. Theoretical modelling strongly suggested that this occurs *via* a complex, but defined, network of interactions^11,12^. Experimental studies concluded that the initial horizontal orientation along the mid-flank of the first iridophores determines the orientation of the horizontally-striped pattern^1,13,14^. However, our initial modelling indicated this might not always be sufficient^12^. Marked dorso-ventral (DV) asymmetries also remain unexplained.

During pattern development, zebrafish grow up to 7.5 fold in length^15^. The growth rate influences the success of stripe formation in some mutants^16^. There is some correlation between *Danio* species pigment and growth patterns, e.g. species with vertical disruptions in the stripes have deeper bodies and a prolonged period of juvenile growth^17^. Building on our model of zebrafish pattern development^12^, we show that growth rates are key determinants of patterning, readily over-riding initial iridophore patterns. By changing growth parameters, whilst keeping pigment cell interactions and initial conditions unchanged, we were able to generate diverse patterns resembling those of other species within the *Danio* genus. Furthermore, altering the degree of asymmetry of growth allowed us to reproduce the DV asymmetry of *Danio* patterns, leading us to predict that DV growth in *Danio rerio* is biased towards ventral regions, and may also explain the previously unexplored DV asymmetry of *Danio nigrofasciatus*.

## Results

### Initial iridophore distribution is not predictive of the orientation of pattern

To test the hypothesis that orientation of the initial iridophore stripe determines the horizontal orientation of the final pigment pattern in zebrafish, we performed an *in silico* experiment using our agent-based model to simulate development up to the J+ stage^12,15^. As demonstrated previously^12^, with the initial condition set to include 3 rows of iridophores horizontally across the centre of the domain as seen in WT fish, we consistently observe horizontally-organised stripes at the end of simulations with WT growth parameters (Fig. 1a and Fig. 1a’, Supplementary Movie 1). Surprisingly, even when the initial position of the iridophores in our simulation was dramatically altered to a vertical bar, a cross-shape or a central square (whilst all other conditions remained unchanged), the simulated patterns consistently showed distinct horizontal stripe elements in the final orientation of the pattern (Fig. 1b-1d, Fig. 1b’-d’ and Supplementary Movies 2-4). To provide an objective quantification of striping, we used a pair correlation function (PCF) to determine and quantify spatial correlations. We took the mean (over 50 repeats) of the final pattern PCF^18^ in the horizontal (and separately vertical) direction, for each of the initial conditions. Regardless of the initial iridophore distribution, there was strong pairwise correlation in the vertical direction, but not the horizontal (Fig. 1a’’-1d’’), showing that the orientation of the patterns formed at stage J+ were dominated by horizontal elements, directly contradicting the hypothesis that the initial interstripe orientation plays the defining role in pattern orientation.

**Figure 1.**
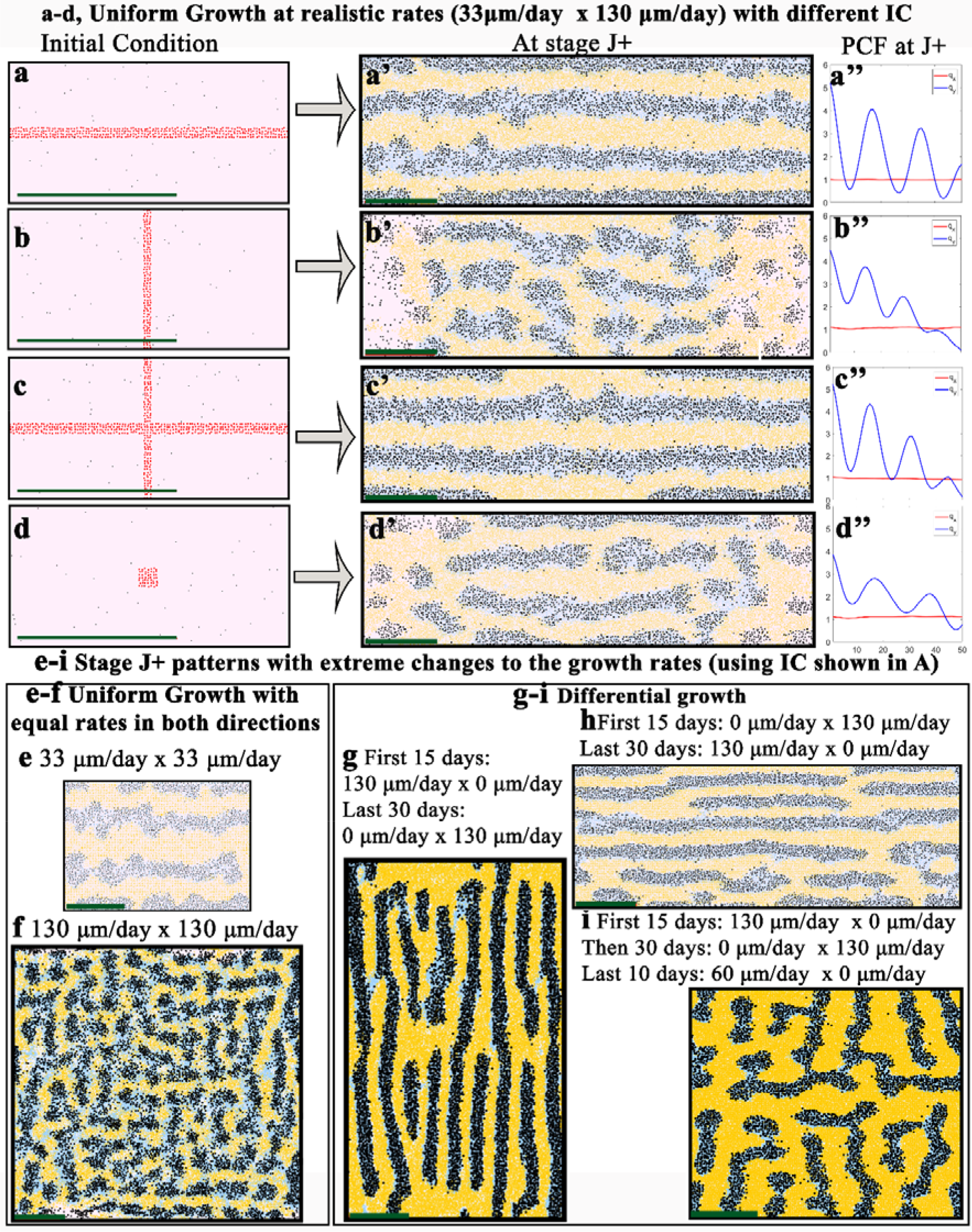
Pigment pattern orientation depends on growth, not just orientation of initial interstripe. **a-d**, Hypothetical initial conditions for the position of dense iridophores (coloured in red for clarity). **a**, WT *Danio rerio* initial conditions. **b**, Vertical stripe of iridophores. **c**, Cross shape of iridophores. **d**, Central square of iridophores. **a’-d’**, Predicted pattern at stage J+ given respective initial conditions and normal growth rates: 33μm/day in the vertical direction and 130μm/day in the horizontal direction. **a’’-d’’**, Mean PCF of 50 repeats of stage J+ simulations with initial condition (**a-d**) in the horizontal axis (red) and vertical axis (blue). **e-i**, Simulated patterns at stage J+ with WT *Danio rerio* initial conditions (**a**) but different rates of growth, as indicated. The green scale bar indicates 1mm in all figures. Alternative simulations to those represented in **a-i** are given in Supplementary Movies 1-9.

### Changing the rate of growth markedly changes patterning

We observed that horizontal pattern elements consistently correlate with domain growth, and, in particular, faster rostrocaudal growth relative to dorsoventral growth (Supplementary Movie 2), suggesting that the dominance of horizontal pattern elements may reflect a determining role of differential growth in pattern formation. To test this hypothesis, we performed simulations in which the initial iridophore interstripe was maintained horizontally, but the growth rates in horizontal and vertical axes were altered (Supplementary Movies 5-9). First, we set the growth rate to be equal in both directions; when the growth rate was slow stripes were maintained (Fig. 1e; Supplementary Movie 5), but when growth was faster, the pattern degenerated into a broken labyrinthine configuration (Fig. 1f; Supplementary Movie 6). We then explored the impact of time-dependent growth rates. Setting dorsoventral growth rates to 130μm per day (and 0μm per day rostrocaudally) for the first 15 days and then 130μm per day rostrocaudally (and 0μm per day dorsoventrally) for the next 30 days generated a pattern of fine vertical bars (Fig. 1g; Supplementary Movie 7), with a switch from early horizontal striping to increasingly vertical barring triggered by the changed growth orientation (Supplementary Movie 7). In contrast, inverting this growth pattern, generated a maze-like pattern (Fig. 1h; Supplementary Movie 8): here transient horizontal stripes are overwhelmed by dorsoventral growth, rapidly becoming a vertically-barred pattern, before modification to form extensive horizontal stripe elements upon the switch to rostrocaudal growth (Supplementary Movie 8). When simulated growth alternated multiple times between being exclusively rostrocaudal and dorsoventral, patterns became labyrinthine with strong horizontal and vertical stripe elements, again driven by the changing growth orientations (Fig. 1i; Supplementary Movie 9). Thus, growth rates are likely to play a key role in controlling major pattern features.

### Impact of biased growth on multiple features of zebrafish pigment pattern

We hypothesised that *asymmetric* growth in the DV axis might explain the apparent dorsal shift of the X0 interstripe during metamorphosis^19^, and the relative width of secondary stripe formation. To assess asymmetry in body growth during pigment pattern formation, we repeatedly measured the relative dorso-ventral position (RDVP) of the centre of the X0 interstripe of eight individual zebrafish. We found that the RDVP of the centre of X0 moves from relative position 0.5±0.01 at stage PB (8mm SL), to relative position 0.4±0.01 by stage J+ (16 mm SL, shown qualitatively in Fig. 2a; rate of translocation quantified in Fig. 2b). During pattern metamorphosis, stripes and interstripes form sequentially from the central interstripe. Assuming symmetric growth about the DV axis (and all cell-cell interactions are symmetric), we would expect that stripes 2D and 2V would appear at the same time. We found that whilst in all cases stripes 1D and 1V both became visible simultaneously (mean±sd: 9.69 ± 0.45 mm SL), stripe 2V always appeared before stripe 2D (2V; 11.81±0.84 mm SL, 2D; 14.69±1.16 mm SL, Fig. 2c). This difference is most readily observed when assessing development of the stripes close to the caudal fin (Fig. 2d). At 10 mm SL, the domain comprises the two stripes 1D and 1V at the very dorsal and ventral regions, respectively, separated by the X0 interstripe. As the fish grows, whilst stripe 1D remains at the very dorsal region of the fish, stripe 1V relocates dorsally over time. By 16 mm SL the 1V stripe is close to being aligned with the central caudal fin stripe, allowing space for a new 2V stripe to appear ventrally. Meanwhile 1D remains at the very dorsal part of the domain. These observations suggest that growth might be ventrally-biased in the DV axis. Implementing our model^12^ with unbiased growth (Fig. 2e, upper panel) generates a poor match to the measured X0 position, since the simulated X0 stripe remains in the middle of the domain (RDVP 0.5). We adapt the model so that the domain grows more ventrally than dorsally (see Supplementary Material). Using the equation relating the position of X0 and the standard length measured from eight fish (derived using linear regression), we determine parameters for biased growth that give a good fit to the real data based on the position of X0 over time (see Supplementary Material). The impact of this bias on X0 position is displayed in Fig. 2e (lower panel). The position of X0 migrates dorsally in our simulations as the fish grows, at a rate similar to that quantified in real fish.

**Figure 2:**
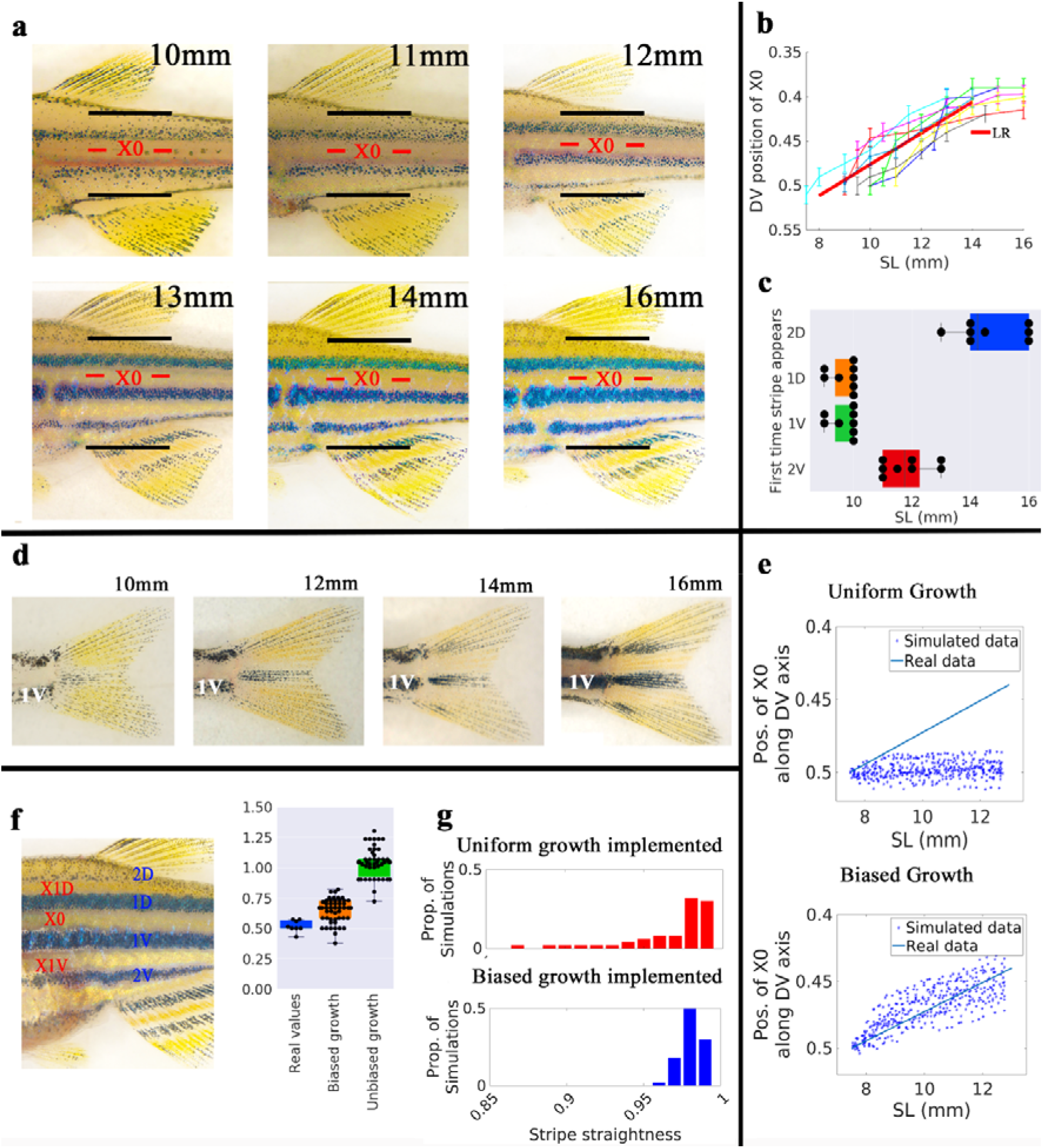
Dorsal repositioning of the first (X0) interstripe can be explained by ventrally-biased growth during pattern development. **a**, WT pattern development on the body. Black lines indicate dorsal and ventral extremes of posterior trunk, red line the DV midline of the X0 interstripe. **b**, Relative position of X0 along the DV axis against standard length (SL) for each fish (N=8). Thin coloured lines plot measured values of each fish, error bars show precision measures (approx.. +10%); thick red line represents the results of undertaking a linear regression (LR) of all of the points (see Methods). **c**, Boxplot showing temporal appearance of stripes during development indicating SL at time when each stripe first visible. Lines indicate, in order, the minimum value, lower quartile, median, upper quartile and maximum value. Stripes 1D and 1V always first became visible on the same day. **d**, Pattern development on the body close to the caudal fin. Stripe 1V is initially adjacent to the most ventral part of the fish at 10 mm SL, but has shifted centrally by 16 mm SL. **e**, Comparison of the variation of the simulated position of X0 with SL in WT fish with unbiased (upper) and biased growth (lower). The blue line indicates the line of best fit for the real data and is the same as the red line shown in **b**. **f**, Juvenile fish at stage J+ (13 mm SL) to show relative size of stripes and interstripes. Graph shows box plot comparison of the ratio of DV extent of stripes 2D to 2V with real data and 50 simulations of WT fish at stage J+ using biased and unbiased growth parameters. A box plot of the ratio of 2D to 2V with real data and 50 simulations of WT fish at stage J+ using biased and unbiased growth parameters. **g**, Comparison of straightness of X0 of WT simulations at stage J+ using biased and unbiased growth parameters.

One further limitation of our previous unbiased simulations was the failure to predict the dorsoventral asymmetries in stripe width^12^. We now show that this is corrected by implementing ventrally-biased growth. In real fish, while stripes 1D and 1V are of approximately equal width, stripe 2D is 0.52±0.05 times as thick as stripe 2V. We propose that this results from biased growth, increasing the space available for stripe 2V relative to stripe 2D. Comparing the ratio of the widths of 2D to 2V in real fish, with those seen in simulations employing either unbiased or biased growth (Fig. 2f) shows that using unbiased growth, the mean stripe width ratio of 2D and 2V is, unsurprisingly, 1±0.02, which does not match the real data. In contrast, implementing biased growth the mean stripe width ratio is 0.58±0.05, similar to the real value. Interestingly, we found that biasing the growth ventrally also increased the mean straightness of the X0 interstripe, as well as reducing the variance (biased growth: 98±0.08%, unbiased growth: 96.9±2.6%, Fig. 2g).

Finally, we investigate ventrally-biased growth as an alternative explanation for observed changes in melanophore differentiation and location during metamorphosis. Parichy and Turner document the initial RDVP of melanophores observed over the course of development (Fig. 3a)^16^. They showed that melanophore births which are initially uniformly distributed between 0-0.25 RDVP and 0.75-1 RDVP, become biased ventrally during pattern metamorphosis. We predicted that this observation might be a consequence of ventrally-biased growth. Consistent with this, the position of melanophore differentiation in simulations using our model with unbiased growth do not match the published observations; instead, melanophore differentiation is symmetric along the DV axis (Fig. 3a’). In contrast, when we incorporate biased growth (Fig. 3a’’)) we observe the same ventral bias as documented in real fish (Fig. 3a), reflecting the dorsal shift of the X0 stripe.

**Figure 3.**
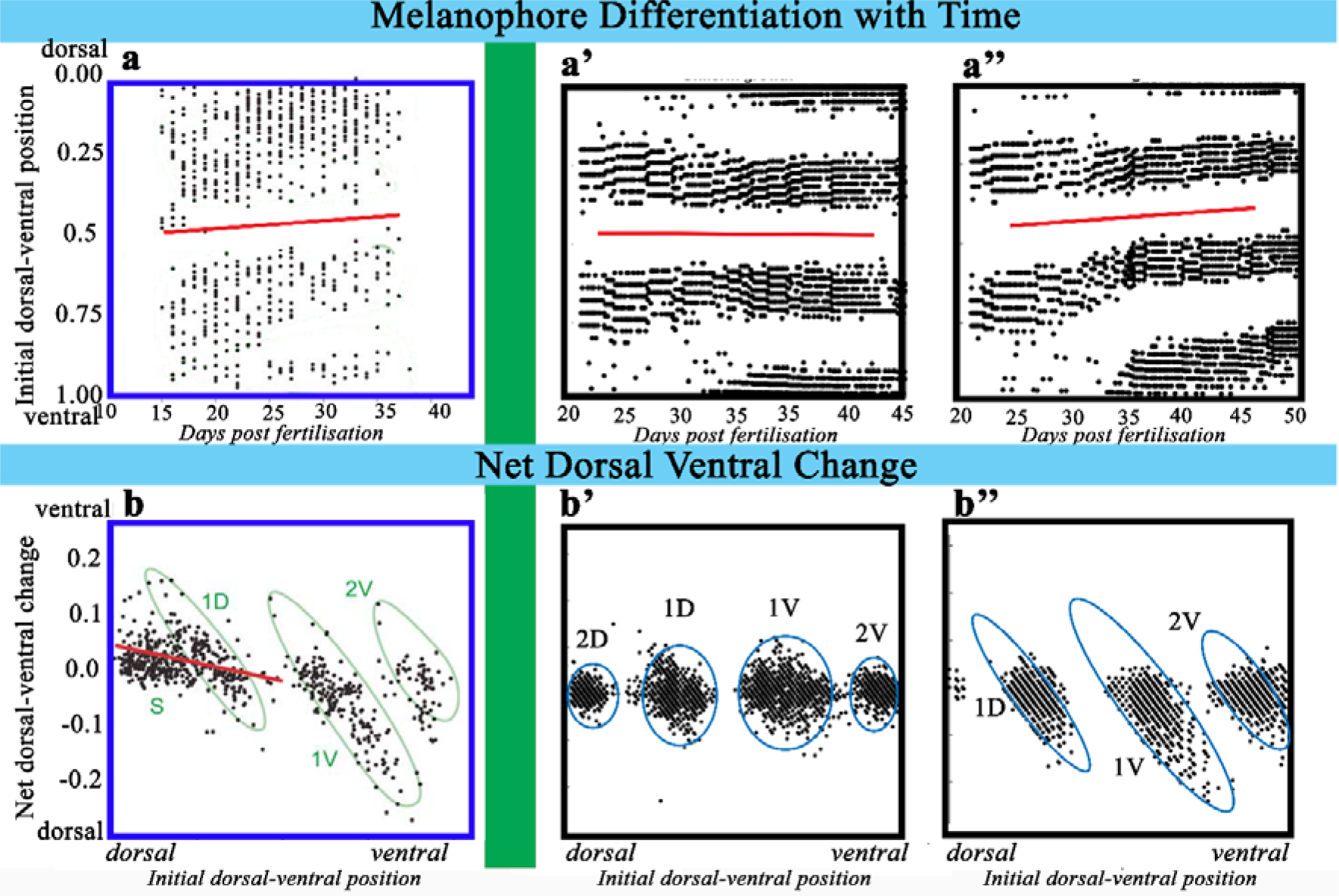
Melanophore births (upper) and movements (lower) in WT fish and simulations of WT fish with and without biased growth. **a**, Melanophore differentiation over time as documented by ^16^. Shown are initial RDVPs of melanophores first appearing on the day of imaging (*x*-axis). Red line indicates predicted centre of the X0 interstripe. **a’-a’’**, RDVP of melanophores at time of birth for representative WT simulations with (**a’**) unbiased growth, (**a’’**) biased growth. Note that the data in **a** includes scale melanophores present in the very dorsal region between stripes 1D and 2D, which are not included in our model, making the differentiation of melanophores in stripes 1D and 1V cleaner in our simulations (**a’-a’’**) than in the experimental data (**a**). **b**, Net changes in RDVPs of melanophores (*y*-axes) plotted against the initial dorsal–ventral positions at which these cells were first identified (*x*-axes) as documented by ^32^. Red line is drawn for melanophores that appeared between 21 and 26 dpf computed using linear regression and includes embryonic melanophores which are not explicitly incorporated in our model. Ellipses are drawn around cells that ultimately contributed to dorsal and scale melanophores (S), the adult primary dorsal melanophore stripe (1D), the adult primary ventral melanophore stripe (1V) and the first adult secondary melanophore stripe at the ventral margin of the flank (2V). **b’-b’’**, Net changes in dorso–ventral positions of melanophores (*y*-axes) plotted against initial dorso–ventral positions at which melanophores first appear (*x*-axes) over a representative WT simulation with (**b’**) unbiased growth and (**b’’**) biased growth. Since net changes are calculated by subtracting the starting position from the final position for each melanophore, negative net changes represent dorsal movements. Panels **a** and **b** were reproduced from ^16^ and licensed under CC-BY 3.0 (http://creativecommons.org/licenses/by/3.0); published by The Company of Biologists Ltd.

Parichy and Turner also document the net change in RDVP of melanophores from their RDVPs at the time of initial differentiation^16^ (Fig. 3b), observing that melanophores tended to relocate dorsally during pattern development and that this was most prominent for the melanophores comprising stripe 1V. They interpreted this change as resulting from dorsally-biased migration. Whilst this could undoubtedly be achieved through a complex mechanism specifying biased migration, a more parsimonious explanation might be biased growth. Model simulations with unbiased growth display limited and dorsoventrally-symmetric net changes in RDVP of melanophores during growth; these data only poorly match the original observations (Fig. 3b’, compare Fig. 3b)). However, when we incorporate ventrally-biased growth, we observe larger shifts in RDVP, with clear dorsal biases, which are most prominent in stripe 1V (Fig. 3b’’), matching closely the key features of the original observations in real fish. We conclude that ventrally-biased growth is sufficient to explain the observed relocations of melanophores during metamorphic growth, suggesting that they likely do not result from biased melanophore migration as such.

### Biased growth explains asymmetry in ‘missing cell-type’ and *rose* mutants

Pigment pattern DV asymmetry is also observed in ‘missing cell-type’ mutant zebrafish, i.e. *pfeffer (pfe)*, *shady (shd)* and *nacre (nac)* which lack xanthophores, iridophores and melanophores respectively. These asymmetries are most prominently observed in adulthood and not in juvenile stages^1^. To assess whether these may also be explained by ventrally-biased growth, we adapt the bias parameters in our model so that the expected position of X0 at adulthood (RDVP 0.38) is reached by the end of our simulation window at 13.5 mm SL. This acceleration is necessary to overcome the current limit imposed on the duration of our model simulations (terminating in juvenile stages (13.5mm SL)) by a lack information about L-iridophores, which are postulated to play a role in pattern maintenance in these later metamorphic stages^1,20^. Further discussion and motivation for this adaptation is given in the supplementary material.

We begin by showing that implementing this modified bias does not affect the model’s recapitulation of the expected WT pattern (Fig.4a-d). Implementing the bias in each of the “missing cell-type” mutants reveals that biased growth introduces asymmetries to the patterns in a manner qualitatively matching those observed in real fish (Fig. 4e-p). For example, in Fig. 4h, biased growth within our model results in simulated *pfe* mutant patterns with spot-like features in ventral ‘stripes’, contrasting with more connected spots, forming stripe-like features, dorsally. In Fig. 4l, biased growth results in simulated *shd* mutant patterns with ventral melanophore spots being spaced out and rounded, whereas dorsal melanophore spots are more closely spaced, as in real *shd*. In Fig. 4p we are able to replicate the expanded X0 interstripe region and bleb-like extensions of dense iridophores and xanthophores seen in real *nac* (Fig. 4p rectangle and Fig. 4n).

**Figure 4.**
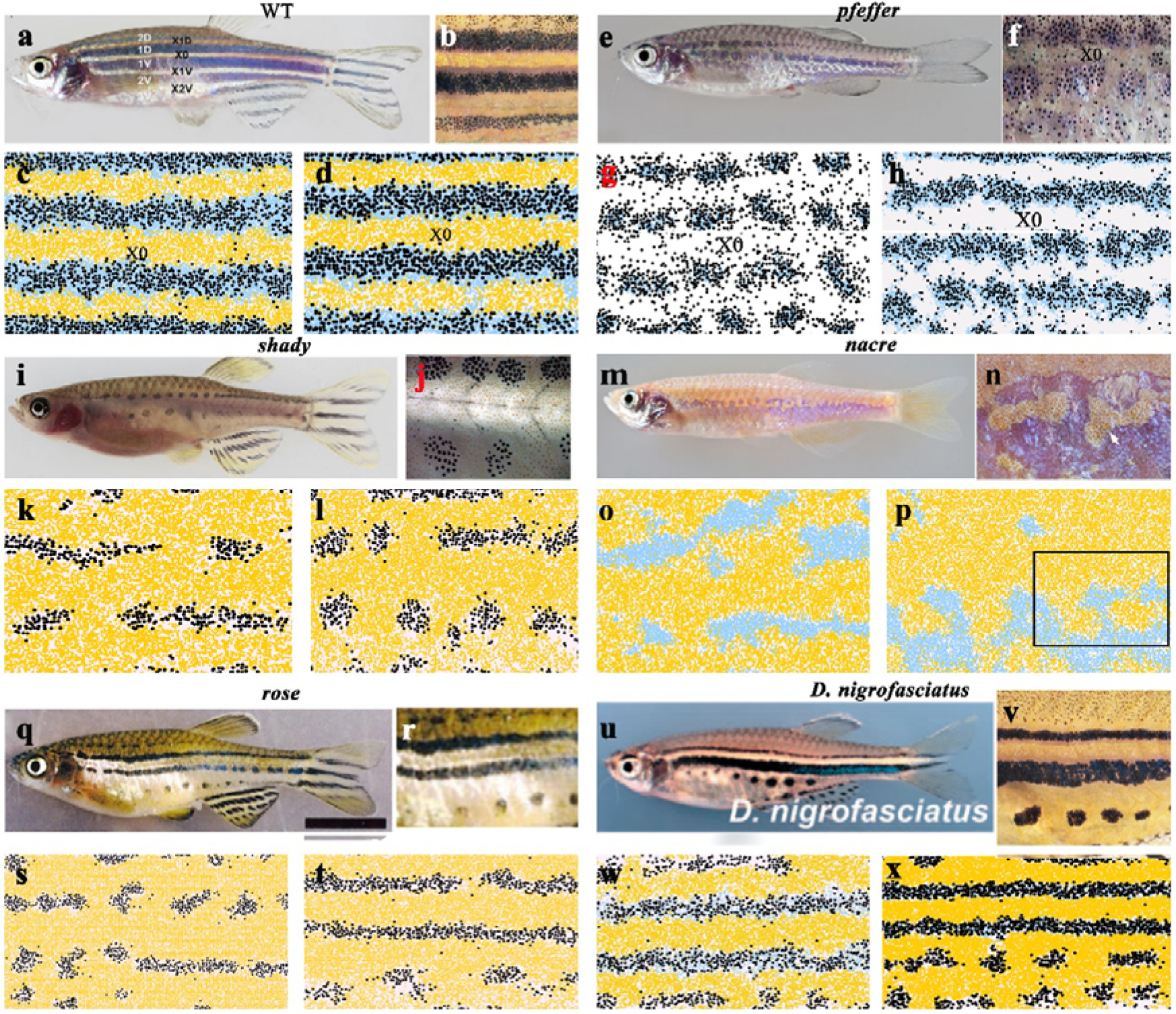
Ventrally-biased growth improves pattern predictions for WT and *pfe, shd, nac* and *rse* mutants and for *Danio nigrofasciatus*. **a,e,i,m,q,u** Images of adult WT*, pfe, shd, nac, rse* and *Danio nigrofasciatus* fish respectively. **b,f,j,n,r,v** Close-ups of the adult patterns displayed by WT, *pfe, shd, nac, rse* and *Danio nigrofasciatus* respectively. **c,g,k,o,s,w** and **d,h,l,p,t,x** Representative simulated patterns of adult WT*, pfe, shd, nac, rse* and *Danio nigrofasciatus* at stage J+ with unbiased and ventrally-biased growth respectively. In each case, simulations performed with biased growth provide better qualitative agreement with the real adult fish than simulations performed with unbiased growth. The box in **p** highlights a region with bleb-like pattern elements repeatedly observed in the simulated *nac* pattern, and which matches blebs observed in real fish, as shown in **n**. Images **a,b,e,f,l,j,m,n,u,v** and **q,r** are reproduced from ^1^, ^22^ and ^43^ respectively and are licensed under CC-BY 3.0 (http://creativecommons.org/licenses/by/3.0); published by The Company of Biologists Ltd.

A notable DV pigment pattern asymmetry is seen in *rse* mutants, in which a reduction in iridophore proliferation generates a partially spotted pattern, with stripes dorsal and ventral to the X0 interstripe, but with more ventral stripes broken into spots. When we reduce iridophore proliferation in simulations to mimic the known biology of *rse* mutants with unbiased growth we generate a symmetric ‘*rse*’-like pattern^12^: two stripes (often broken into spots) symmetric about the centre of the domain (Fig. 4s). However, adding biased growth replicates the real, dorsoventrally asymmetric *rse* phenotype (Fig. 4t).

### Biased growth may contribute to the DV asymmetry of *Danio nigrofasciatus*

*Danio nigrofasciatus*, a close relative of zebrafish, displays two stripes dorsally and small melanophore spots ventrally (Fig. 4u,v). We interpret these features as resulting from evolutionary alterations of the zebrafish pigment patterning mechanism as described by ^21,22^ (see Methods). Implementing these changes, but without biased growth, our model predicts a DV-symmetric pattern of four horizontal stripes, with 2D and 2V being broken (Fig. 4w). Upon incorporating ventrally-biased growth, we reliably (100/100 simulations) recapitulate the asymmetry of the *D. nigrofasciatus* pattern in our simulations (Fig. 4x).

## Discussion

Here we demonstrate an unexpectedly important role for growth in determining zebrafish body patterns. The importance of growth in influencing development and the final pattern has been suggested theoretically^23-30^ and correlated with inter-specific pattern differences^17^, but these have not been tested in the context of real biological data, beyond the effects on specific pattern details such as stripe width^31^.

The results of our modelling allow us to reject previous hypotheses that suggested that the iridophore stripe acts as a crucial pre-pattern driving the final pattern orientation. In particular, we show that even drastic changes to the initial iridophore positioning, such as to a vertical stripe, yields relatively small changes to the final pattern. In contrast, dramatic changes in pattern (to maze-like and vertically barred patterns) can be induced by altering the timing and rate of differential growth.

We provide evidence that growth may be biased ventrally in zebrafish by showing *in silico* how it can substantially improve multiple previously unresolved inaccuracies of simulations of both WT and mutant patterns. For example, we show that growth bias in WT *Danio rerio* provides novel explanations of the dorsal translocation of X0 during development, previously attributed to melanophore differentiation and migration^32^. Growth asymmetry also explains the ratio of the stripe width of 2D to 2V, as well as improving the overall straightness of X0. The same phenomenon also explains the asymmetries observed in multiple *Danio* pigment pattern mutants. *D. nigrofasciatus* also has a striking DV asymmetry to its pattern; combining the proposed patterning mechanism of *D. nigrofasciatus* (as documented by ^21,22^) with ventrally-biased growth, we are able to generate its characteristic pattern. Further work is required to determine why we do not generate a 1V stripe thicker than the 1D stripe as seen in *D. nigrofasciatus*; the real pattern may result from persistence of embryonic melanophores, which play an important role in *D. nigrofasciatus*^21^, but are not explicitly incorporated in our model.

A number of factors likely explain growth’s prominent role in determining pattern. Firstly, skin growth creates space for cell differentiation, as seen in growing angelfish (*Pomacanthus*), which generate new stripes between existing stripes as the developing skin expands^33^. Growth that is biased allows more space for differentiation in some regions, whilst limiting space in others; this likely underlies the asynchronous appearance of stripes 2D and 2V. Secondly, growth can also interrupt cellular interactions by dispersing cells. We suggest that ventrally-biased growth produces asymmetry by restricting space for new cells dorsally, thus maintaining the stripes in those regions, whilst creating space ventrally, reducing the effectiveness of cellular interactions and increasing the potential for stripes to break up into spots in those regions. Thirdly, growth can affect the directionality of the pattern by spreading cells along a particular axis, increasing the probability that a pattern becomes biased.

There are some limitations to our approach, principally due to the current poor understanding of the role of the late appearing L-iridophores in pigment pattern maintenance^20^; in future, detailed characterisation of the patterning role of L-iridophores could be incorporated into our model to allow a more complete understanding of pigment pattern formation. A further limitation is that the model does not currently include the sites of entry of melanophores onto the flank nor any early differentiating and highly migratory cells, so that we cannot entirely rule out a role for migrating melanophores in contributing to dorsoventral asymmetry.

Nevertheless, our results emphasise the importance of growth and, especially differential growth, of the skin field in determining pigment pattern; growth position, growth rate, and temporal changes in each need to be considered when investigating pattern development of *Danio*, other fish and vertebrates more generally. Furthermore, such skin growth needs to be considered in reference to the proliferation (cell division) rate of the pigment cells themselves. These mechanisms are likely to be critical in other contexts, including the development of varied pigment patterns within human pathophysiological skin conditions, such as piebaldism^34,35^, where they will act alongside the well-known impact of clonal expansion of pigment cells.

## Methods

### Ethics Statement

This study was performed with the approval of the University of Bath ethics committee and in full accordance with the Animals (Scientific Procedures) Act 1986, under Home Office Project Licenses 30/2937 and P87C67227.

### Repeated imaging of juvenile fish

We repeatedly imaged N=8 fish every 2 days from 23 dpf for a total of 6 repeats using a Zeiss AxioZoom V16 with Axiocam 506 color camera and Zen Blue software. Fish were first anaesthetised in a bath of 0.085% 2-phenoxyethanol. Once they were non-touch responsive, the fish were removed from the bath and placed in a shallow pool of 0.085% 2-phenoxyethanol (diluted with system water) and placed under the microscope for imaging. Fish were never anaesthetised for more than 10 minutes. At the end of the experiment, fish were euthanised by Schedule 1 killing.

### Mathematical model of zebrafish pigment pattern formation

#### Mathematical model

The model used for this experiment is explained in detail in ^12^ and we summarise its main features here. The model is a comprehensive on-lattice stochastic mathematical model of zebrafish stripe formation, spanning the developmental period defined by 8 – 13.5 mm SL. The model consists of three stacked lattices representing the three pigment cell layers as observed in the literature^20^: the melanophore layer accommodating melanophores, an iridophore layer which accommodates loose and dense iridophores and a xanthophore layer which accommodates xanthophores and xanthoblasts. The five pigment cell types are modelled as individual agents which occupy lattice sites on these domains. Volume exclusion rules apply so that no site can be occupied by more than one cell. Cells interact, differentiate, proliferate, die and move at a rate consistent with the biological data (provided this is known). The domain grows over time through the insertion of lattice sites at a rate which mimics the natural growth of fish embryos. Boundary conditions are periodic across the horizontal boundaries and reflecting along the vertical boundaries. We implement periodic boundary conditions along the horizontal axis based on the assumption that the rate at which cells leave along these boundaries is approximately equal to the rate at which cells enter the domain at the opposite side. In practice the choice of reflecting or periodic boundary conditions makes very little difference to the appearance of the pattern. The model is updated according to the Gillespie algorithm^36^. To do this we treat all the potential actions, for example cell birth or domain growth as individual ‘events’, each with an exponentially distributed waiting time that corresponds to their mean rate of occurrence. To update the model at any given time *t* = *T*, an exponentially distributed waiting time is generated until the next possible ‘event’ occurs (based on the rates of all of the possible events). Next a uniformly distributed random number determines which event occurs based on the relative probability of each event. For some events to occur there are also additional constraints that must be met (for example, if a cell is attempting to move, the space into which it is attempting to move must be unoccupied). If conditions required for that event to occur are met, the event is implemented, whereas if they are not, then there is no change. Once an event is accepted, the domain is updated accordingly, and time is also updated by drawing an appropriate exponentially distributed random number. This process repeats until we reach the end of pattern metamorphosis marked by the domain reaching a length of 13.5 mm SL. Previously, we have shown that our model can not only consistently replicate qualitative and quantitative features of WT patterns, but also other zebrafish mutant patterns, demonstrating its validity in investigation of zebrafish pattern formation^12^.

#### Implementing growth bias

In our original model when a growth event in the vertical (without loss of generality) direction is chosen to occur, a site for the growth event is chosen uniformly at random from each column^12^. Supposing the vertical extent of the domain is currently *L* sites, uniform unbiased growth dictates that the probability that a growth event occurs at a particular site *i* is 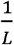. A new site is inserted below the chosen site with probability 0.5, and above the chosen site with probability 0.5. The corresponding sites in that column are shifted upwards to make room for the new site. We incorporate growth bias by changing the probability that a growth event occurs at a given position, such that it occurs with probability *q* below *γL*, where *q* ∈ [0,1] and *γ* ∈ [0,1]. We determined the values of the ‘bias parameters’ *q* and *γ* for zebrafish by comparing simulated domains against five experimentally measured characteristics described in the supplementary material. Explicitly, we found that bias parameters *q* = 0.28, *γ* = 0.15 replicate normal WT growth. In the later sections we employ a stronger bias *q* = 0.5, *γ* = 0.25 to bring about the same shift in X0 at an earlier time point (see supplementary material).

#### Position of X0

The position of X0 is calculated by first compiling the set of positions *T*_*i*_ and *B*_*i*_ that define the maximum (*T*) and minimum (*B*) extent of the X0 interstripe respectively in each column, *i*. In this way 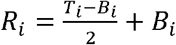, the set of positions centred within the X0 interstripe, is a one-dimensional representation of the central interstripe. (The full algorithm for generating *R*_*i*_ is also used to determine the tortuosity of the stripes and is given in detail within the supplementary material of the paper: ^12^) The position of X0 in terms of lattice sites is the mean of *R*_*i*_ which we define as *p*. The relative position of X0 is then calculated as:

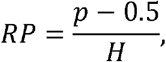

where *H* is the height of the domain.

#### Stripe straightness

To measure stripe straightness, we first generate a one-dimensional representation (*R*_*i*_) of the stripe (as described in the above section for the central interstripe). From this line *R*_*i*_ we calculate the stripe straightness *SS(x)*, measured as the ratio of the total length of our line represented by the set of points *R*_*i*_, which we refer to as *R*, to the straight-line distance between its ends (*C*).

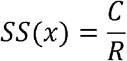

For more detail see ^12^.

#### Implementation of *Danio rerio* mutants

##### shd

The gene *shady* (*shd*) encodes zebrafish *leukocyte tyrosine kinase* (Ltk) which plays a role in S-iridophore specification ^37-39^;. As a result, strong *shd* mutants lack iridophores. To simulate this defect, we remove all S-iridophores from the initial domain. New S-iridophores are only generated by the proliferation of existing S-iridophores, thus this removes the generation of any S-iridophores throughout the simulation.

##### nac

The gene nacre (*nac*) encodes transcription factor Mitfa^40^. *nac* mutants lack melanocytes throughout embryonic and larval development. To simulate this, we remove all melanocytes from the initial conditions. We set the rate of new melanophore differentiation to be zero.

##### pfe

Gene *pfeffer* (*pfe*) encodes for *colony stimulating factor 1 receptor* (*csf1ra*) which plays a role in xanthophore specification and migration^41,42^. In strong alleles, adult fish exhibit no detectable xanthophores in the body of embryos. To simulate this, we remove all xanthophores and xanthoblasts from the initial domain. Since new xanthophores are only generated by the proliferation of existing xanthophores and the differentiation of xanthoblasts, this means that no xanthophores appear on the domain during the simulation.

##### rse

Rose (*rse*), encodes the Endothelin receptor B1a and has been shown to acts cell-autonomously in S-iridophores; homozygous mutants result in a reduction of S-iridophores to approximately 20% of that seen in WT (observed in stage PB and adult fish). To simulate *rse* we change the initial conditions so that the number of iridophores on the domain is one fifth of the usual number. We place the remaining iridophores uniformly at random within the central three rows, since some S-iridophores still appear along the horizontal myoseptum in *rse*. We also reduce their rate of proliferation to one fifth of the usual number.

For more explicit detail about the simulation these mutants, please see the supplementary material of ^12^.

#### Implementation of *Danio nigrofasciatus*

A brief summary of the known developmental differences between *Danio nigrofasciatus* and *Danio rerio* is given by ^21,22^: during pattern metamorphosis, fewer metamorphic melanophores differentiate, and instead a greater number of embryonic melanophores survive. Iridophores do not proliferate as frequently. We interpret this in our model as a decreased rate of differentiation of melanophores in later stages; increased survival rate of melanophores in early stages (as a proxy for the number of embryonic melanophores that survive but are not represented explicitly in our model) and a decreased rate of iridophore proliferation.

#### Dorsal -ventral relocation of melanophores

To determine the dorso-ventral relocation of melanophores we recorded the relative DV position of each melanophore upon appearance and subtracted its final position to determine its net movement.

#### Pair correlation function

The pair correlation function (PCF) used in Fig. 1 is as described by ^18^. We use the PCF on the melanophore domain. The value of the PCF at any given distance *d* in the horizontal or vertical direction respectively is the number of melanophores on the domain that are exactly this distance apart divided by the number we would expect if the same number of melanophores were places on the domain uniformly at random. If the value of the PCF at a certain distance is unity this suggests that there is no spatial correlation between cells at that distance. If the value of the PCF is greater than unity then this suggests there is spatial correlation at this distance. If the value of PCF is less than unity then this suggests that there is spatial anticorrelation at this distance.

### Statistics

#### Linear regression

To create the line of best fit in Fig. 2B we performed linear regression using the scikit-learn package in python. We used model type Linear Regression from which we determined the intercept and slope of the line.

## Data availability

Numerical counts data are available from the corresponding author on request.

## Code availability

The pattern model code is available at https://github.com/JenniferOwen/Zebrafish-stripe-model

## Acknowledgements

We acknowledge the technical staff within the Department of Biology & Biochemistry at the University of Bath for technical support & assistance in this work. We thank D. Parichy, A. Perry and J.H.P Dawes for comments on the manuscript. Funding was provided by the BBSRC to J.P.O. (SWBio DTP Studentship) and to R.N.K. (Grant number BB/L00769X/1).

## Author contributions

J.P.O. developed the model, performed the simulations and experiments, analysed the data and drafted and revised the manuscript. C.A.Y. and R.N.K. designed and supervised the study and revised the manuscript.

## Competing interests

The authors declare no competing interests.

## Notes

https://github.com/JenniferOwen/Zebrafish-stripe-model

### Competing Interest Statement

The authors have declared no competing interest.

